# Serotonin signaling in the rat prefrontal cortex is required for Retrieval-Induced Forgetting

**DOI:** 10.1101/2025.03.05.641624

**Authors:** María Belén Zanoni, Mariana Imperatori, Pablo Koss, Beatriz Agustina Ortega, Melisa Bentivegna, Cynthia Katche, Michael C. Anderson, Pedro Bekinschtein, Noelia V. Weisstaub

## Abstract

Forgetting is a ubiquitous phenomenon actively promoted in many species. The act of remembering some experiences can cause forgetting of others in both humans and rats. We previously found that when rats retrieve a memory to guide exploration, it reduces later retention of other competing memories encoded in that environment. As with humans, this retrieval-induced forgetting (RIF) relies on prefrontal control processes, is competition-dependent, and cue-independent. RIF is thought to be driven by inhibitory control signals from the prefrontal cortex that target areas where memories are stored. Serotonin plays a crucial role in behaviors requiring high cognitive demand, including memory processes, partly through its modulation of prefrontal cortex activity. However, its potential involvement in regulating active forgetting remains unexplored. Here, we had rats perform a task known to induce RIF and pharmacologically manipulated the activity and signaling of serotonin receptors 5-HT1A, 5-HT2A, and 5-HT2C in the medial prefrontal cortex (mPFC), as well as to inhibit downstream effectors. Our findings reveal a specific role for prefrontal serotonin signaling in RIF. Whereas 5-HT2C receptor manipulation had no effect, activating 5-HT1A or blocking 5-HT2A receptors in the mPFC abolished RIF. By contrast, activating 5-HT2A receptors promoted RIF under conditions in which it is normally reduced. Further analyses identified the PI3K/AKT pathway as a downstream effector of 5-HT2A receptor signaling, suggesting a specific molecular mechanism through which serotonin modulates inhibitory control over memory. These results uncover a previously unrecognized serotonergic modulation of adaptive forgetting, identifying specific receptor subtypes that link prefrontal serotonin signaling to inhibitory control over memory competition.

**Significance statement:** Forgetting is an essential, active process that helps the brain stay flexible and focused. In both humans and animals, remembering one thing can lead to forgetting competing memories, a phenomenon called retrieval-induced forgetting (RIF). Our research shows that serotonin, a neurotransmitter best known for its role in mood, also actively regulates which memories are forgotten. In rats, we found that serotonin signaling in the prefrontal cortex regulates adaptive forgetting: 5-HT2A receptor signaling promotes RIF, whereas activation of 5-HT1A receptors disrupts it. We further identified PI3K/AKT signaling as a key downstream effector of 5-HT2A receptor activation in this process. These findings matter because the mechanisms of active forgetting remain poorly understood, and their disruption may contribute to psychopathological conditions.

## Introduction

In the face of changing circumstances, facilitating the forgetting of outdated information may be an adaptive strategy (Sangha et al., 2005; Shuai et al., 2010; Benna and Fusi, 2016; Epp et al., 2016). Historically, forgetting was conceptualized as a passive process, arising either from changes in the contextual cues required to retrieve older memories or from the gradual loss of the engrams supporting them. Over the past two decades, however, evidence from flies and rodents has revealed the existence of dedicated active forgetting mechanisms (for reviews, see (Medina, 2018); (Davis and Zhong, 2017)), some of which may account for findings from earlier human studies.

One well-characterized example is retrieval-induced forgetting (RIF): when people or rats recall a past event, competing memories that interfere with retrieval become more likely to be forgotten (for reviews, see Anderson et al., 1994; Anderson and Hulbert, 2021; Marsh and Anderson, 2024). In both species, RIF requires prefrontal engagement during the selective retrieval practice phase (Wu et al., 2014; Bekinschtein et al., 2018; Gallo et al., 2022) and is competition-dependent and cue-independent, meaning that the forgotten memory cannot be recovered using alternative cues(Bekinschtein et al., 2018).

Although several mechanisms have been proposed to explain RIF, most can be grouped under inhibition-based hypotheses (Murayama et al., 2014). According to this view, attempting to recall a particular memory also reactivates related memories that compete for retrieval; these competitors are then inhibited to resolve the interference. Beyond its immediate role, this inhibitory process is thought to have a lasting consequence: by reducing the future accessibility of the inhibited competitor, it optimizes retrieval of the target memory in similar situations, at the cost of forgetting the competing trace. RIF is therefore not a mere byproduct of retrieval but an adaptive mechanism through which selective inhibition shapes the accessibility of memories over time.

Converging evidence identifies the prefrontal cortex (PFC) as the source of the inhibitory control signals underlying RIF (Kuhl et al., 2007; Hanslmayr et al., 2010; Staudigl et al., 2010; Ferreira et al., 2014; Penolazzi et al., 2014). According to the Prefrontal Control Hypothesis (Anderson and Hulbert, 2021), active forgetting (e.g., RIF) originates from top-down control signals mediated by the PFC. In humans, the lateral PFC, particularly on the right side, plays a key role in reducing the accessibility of disruptive memories through inhibitory mechanisms. Yet despite progress in characterizing the behavioral and anatomical features of RIF, the neuromodulatory systems that regulate this prefrontal control remain largely uncharacterized, with dopamine acting on D1 receptors in the mPFC as the only such system identified to date (Gallo et al., 2022)

Among these neuromodulators, serotonin is a particularly strong candidate. Its evolutionarily conserved modulatory role, together with its capacity to exert opposing influences on cognition and behavior, positions it well to regulate the kind of graded, competition-dependent inhibition that RIF requires. Serotonergic fibers from the medial and dorsal raphe nuclei densely innervate the PFC (Molliver, 1987; Hensler, 2006), where three receptor subtypes (5-HT1AR, 5-HT2AR, and 5-HT2CR) are most abundantly expressed and exert distinct effects on prefrontal function (Puig and Gulledge, 2011; Leiser et al., 2015a; Švob Štrac et al., 2016). Through these receptors, serotonin modulates cognitive processes closely related to the control demands of RIF, including response inhibition (Hadamitzky and Koch, 2009; Keeler and Robbins, 2011; Wischhof et al., 2011; Macoveanu et al., 2013) and cognitive flexibility (Boulougouris et al., 2007; Kehagia et al., 2010; Baker et al., 2011; Roberts, 2011), as well as attention and working memory (Passetti et al., 2003). Particularly relevant for the present study, we have shown that the 5-HT2AR in the rat mPFC is required for retrieving object-in-context memories, but not the identity of objects alone (Bekinschtein et al., 2013) suggesting that this receptor contributes to resolving interference during episodic memory retrieval, a process that may extend to the inhibitory control of competing memories in RIF.

In this study, we test whether serotonergic signaling in the mPFC contributes to adaptive forgetting, and if so, through which receptor subtypes and downstream pathways. Using selective pharmacological manipulations during the retrieval practice phase of a rat RIF task, we identify dissociable roles for 5-HT1A, 5-HT2A, and 5-HT2C receptors in the mPFC, and implicate PI3K/AKT signaling as a downstream effector of 5-HT2AR activation. Because disruptions in the balance between remembering and forgetting have been linked to psychiatric conditions including depression, anxiety, and intrusive cognition (Anderson and Hulbert, 2021; Ryan and Frankland, 2022; Berry et al., 2024).

## Results

### Differential involvement of mPFC serotonin receptors in RIF

We first asked which serotonin receptors in the mPFC contribute to RIF. To this end, we pharmacologically manipulated 5-HT1A, 5-HT2A, and 5-HT2C receptors during the retrieval practice phase. Serotonin 1A, 2A and 2C are among the most abundant serotonin receptors expressed in the rat mPFC (Pazos and Palacios, 1985; Pazos et al., 1985; Pompeiano et al., 1992, 1994; Leiser et al., 2015b). These receptors play distinct roles in various cognitive tasks that involve mPFC function, underscoring the serotonergic system’s complex regulation of this structure (Zhang and Stackman, 2015; Bacqué-Cazenave et al., 2020).

Our initial analysis examined the role of the G_i_-coupled 5-HT1AR in RIF. After the encoding phase, rats were randomly assigned to the RP, IC, or TC conditions (Fig. 1A) and received bilateral intra-mPFC infusions of either the selective 5-HT1AR agonist 8-OH-DPAT or vehicle (VEH) before the practice phase, or at the corresponding time point in the TC condition. Based on previous findings (Bekinschtein et al., 2018; Gallo et al., 2022), we expected selective retrieval of one object to impair subsequent memory expression for the competing object, and predicted that this effect would be inversely related to 5-HT1AR activity in the mPFC during retrieval practice.

**Figure 1.**
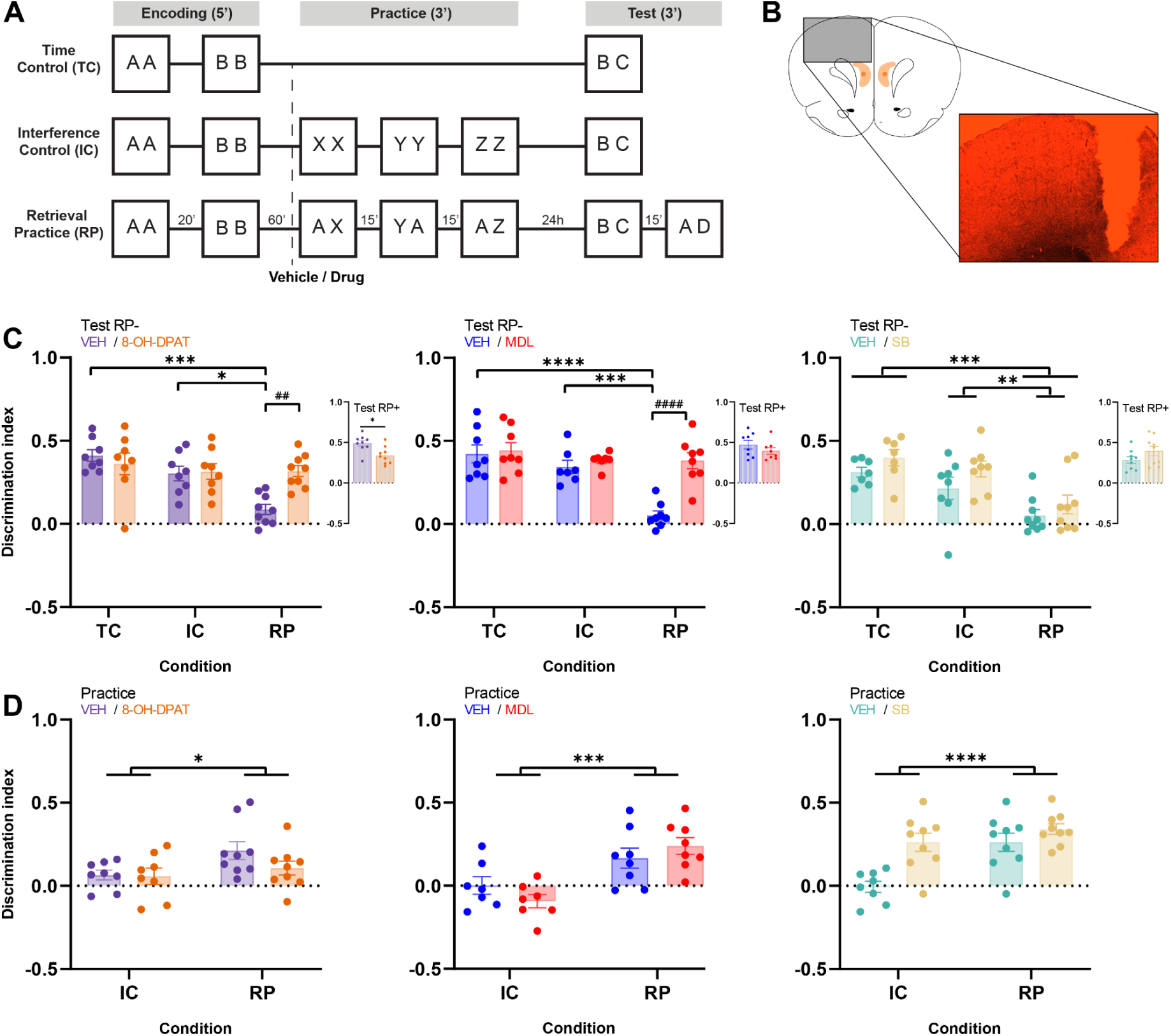
Serotonergic manipulations in the mPFC modulate RIF. **A.** Schematic of the behavioral protocol. Animals were assigned to RP, IC, or TC conditions and performed the task twice, once with drug and once with vehicle (pseudorandomly assigned), in a between-subjects × within-subjects design. The dashed line indicates infusion 15 min before the practice phase. **B.** Schematic of the infusion target and histological section showing the cannula trace. The infusion site lies 1 mm deeper than the cannula tip. **C.** Discrimination index ± SEM at test for three experiments: 5-HT1AR agonist (8-OH-DPAT; n = 25), 5-HT2AR antagonist (MDL11,939; n = 23), and 5-HT2CR antagonist (SB242084; n = 24). In the SB experiment, asterisks indicate the main effect of condition; in the remaining experiments, asterisks (*) indicate significant differences between conditions within VEH and number signs (#) indicate significant differences between pharmacological treatments within conditions (Bonferroni post hoc). Inset: Discrimination index ± SEM for the practiced object. **D.** Discrimination index ± SEM during practice for IC and RP groups across the three experiments. *p < 0.05, **p < 0.01, ***p < 0.001, ****p < 0.0001.

To test this, we analyzed the discrimination index (DI) during the final test phase using a linear mixed model with Condition, Drug, and their interaction as fixed effects, and Subject as a random effect. We found a significant Condition_RP_ x Drug_8-OH-DPAT_ interaction (β = 0.28, _95%_CI [0.10, 0.46], t_(22)_ = 3.26, p < .01). Consistent with RIF, VEH-treated animals showed a lower DI in the RP condition than in the IC and TC conditions ([TC−RP]_VEH_: mean = 0.324, SE = 0.061, t_(22)_ = 5.359, p < .001; [IC−RP]_VEH_: mean = 0.215, SE = 0.061, t_(22)_ = 3.571, p < .05). Importantly, 8-OH-DPAT did not affect performance in the IC or TC conditions (IC: mean_VEH−8-OH-DPAT_ = −0.012, SE = 0.062, t_(22)_ = −0.195, p > .99; TC: mean_VEH− 8-OH-DPAT_ = 0.049, SE = 0.062, t_(22)_ = 0.791, p > .99), but significantly increased DI in the RP condition (mean_VEH−8-OH-DPAT_ = −0.230, SE = 0.059, t_(22)_ = −3.921, p < .01). These results suggest that activation of mPFC 5-HT1AR interferes with forgetting of the competing object (Fig. 1B, left).

We then assessed memory for the practiced object to determine whether this effect was specific to the competitor object. Although 8-OH-DPAT reduced the DI for the practiced object relative to vehicle (β = −0.15, _95%_CI [−0.26, −0.04], t_(8)_ = −3.05, p < .05), both groups still showed significant memory for the practiced object, as indicated by above-zero DI values (one-sample t-test; VEH: t_(8)_ = 14.276, p < .0001; 8-OH-DPAT: t_(8)_ = 9.59, p < .0001). Thus, memory for the practiced object was preserved in both groups (Fig. 1B, left, inset).

To rule out pre-existing differences during retrieval practice that might account for the effects observed at test, we also analyzed exploratory behavior during the practice phase (Fig. 1D, left). As expected from the structure of the protocol, we found a significant main effect of the RP condition on DI across the three practice sessions (β = 0.16, _95%_CI [0.00, 0.32], t_(15)_ = 2.32, p < .05; [IC−RP]: mean = -0.098, SE = 0.045, t_(15)_ = -2.177, p < .05), but no effect of drug. Moreover, 8-OH-DPAT did not alter total exploration time during practice in either the RP or IC groups, indicating that the effect observed at test cannot be readily explained by changes in locomotor, affective, or motivational processes, or by a general impairment in object recognition. Unless otherwise stated, neither condition, drug treatment, nor their interaction affected total exploration time in any analyzed phase.

We next examined the role of 5-HT2A and 5-HT2C receptors, the most abundant members of the 5-HT2 family in the brain. Starting with the 5-HT2AR, animals received local intra-mPFC infusions of vehicle or the selective 5-HT2AR antagonist MDL11939 (MDL) before the practice phase (Fig. 1A).

Twenty-four hours after encoding, we tested memory for the competitor object. As expected, VEH-treated animals in the RP condition showed evidence of RIF, with lower DI values than those in the TC and IC conditions ([TC−RP]_VEH_: mean = 0.370, SE = 0.058, t_(20)_ = 6.346, p < .0001; [IC−RP]_VEH_: mean = 0.291, SE = 0.060, t_(20)_ = 4.826, p < .001). We found a significant Condition_RP_ x Drug_MDL_ interaction (β = 0.31, _95%_CI [0.21, 0.41], t_(20)_ = 6.77, p < .001). Importantly, MDL increased DI only in the RP condition (RP_VEH−MDL_: mean = −0.329, SE = 0.032, t_(20)_ = −10.165, p < .0001), with no effect in the IC (IC_VEH−MDL_: mean = −0.041, SE = 0.035, t_(20)_ = −1.174, p > .05) or TC (TC_VEH−MDL_: mean = −0.019, SE = 0.032, t_(20)_ = −0.590, p > .99) conditions. These results suggest that blockade of mPFC 5-HT2AR interferes with the mechanism underlying RIF (Fig. 1C, center). No treatment differences were observed for memory of the practiced object (Fig. 1C, center, inset).

We also analyzed behavior during the practice phase. Regardless of drug treatment, animals in the RP condition showed higher DI values than those in the IC condition (Condition_RP_: β = 0.16, _95%_CI [4.07e−03, 0.32], t_(13)_ = 2.22, p < .05; [IC−RP]: mean = -0.248, SE = 0.054, t_(13)_ = -4.617, p < .001), indicating greater exploration of the novel object relative to the practiced one (Fig. 1D, center). We also detected a marginal simple effect of condition on the retrieval practice total exploration time (β = 16.43, _95%_CI [0.04, 32.82], t_(13)_ = 2.17, p = .050; [IC−RP]: mean = -14.551, SE = 5.365, t_(13)_ = -2.712, p < .05), but because this effect was independent of drug treatment, it is unlikely to account for the test-phase effect.

We then examined the role of the 5-HT2CR in RIF. To this end, animals received bilateral intra-mPFC infusions of vehicle or the selective 5-HT2CR antagonist SB242084 (SB) before the practice phase (Fig. 1A). In the test phase, we found no significant Condition_RP_ x Drug_SB_ interaction, indicating that the pharmacological treatment did not modify the effect of condition. Regardless of treatment, animals in the RP group showed lower DI values than those in the IC and TC groups, consistent with RIF (Condition_RP_: β = -0.26, _95%_CI [-0.41, -0.11], t_(21)_ = -3.65, p = .001; [TC−RP]: mean = 0.270, SE = 0.054, t_(21)_ = 5.040, p < .001; [IC−RP]: mean = 0.188, SE = 0.052, t_(21)_ = 3.637, p < .01). Likewise, no differences between VEH and SB-treated animals were observed when memory for the practiced object was tested. During the practice phase, SB did not alter the animals’ ability to discriminate between the novel and familiar objects, as reflected by the absence of a drug effect; as expected, animals in the RP condition showed higher DI values than those in the IC condition (Condition_RP_: β = 0.27, _95%_CI [0.11, 0.42], t_(15)_ = 3.73, p < .01; [IC−RP]: mean = -0.336, SE = 0.054, t_(15)_ = -6.236, p < .0001).

### 5-HT2AR activation, but not 5-HT1AR blockade, elicits RIF

We then asked which serotonergic manipulations are sufficient to induce RIF. To do so, we used a modified version of the task that does not normally elicit RIF (Gallo et al., 2022): each pair of objects were presented twice in two consecutive encoding sessions, and the practice phase was separated from the encoding phase by 24 h (Fig. 2A).

**Figure 2.**
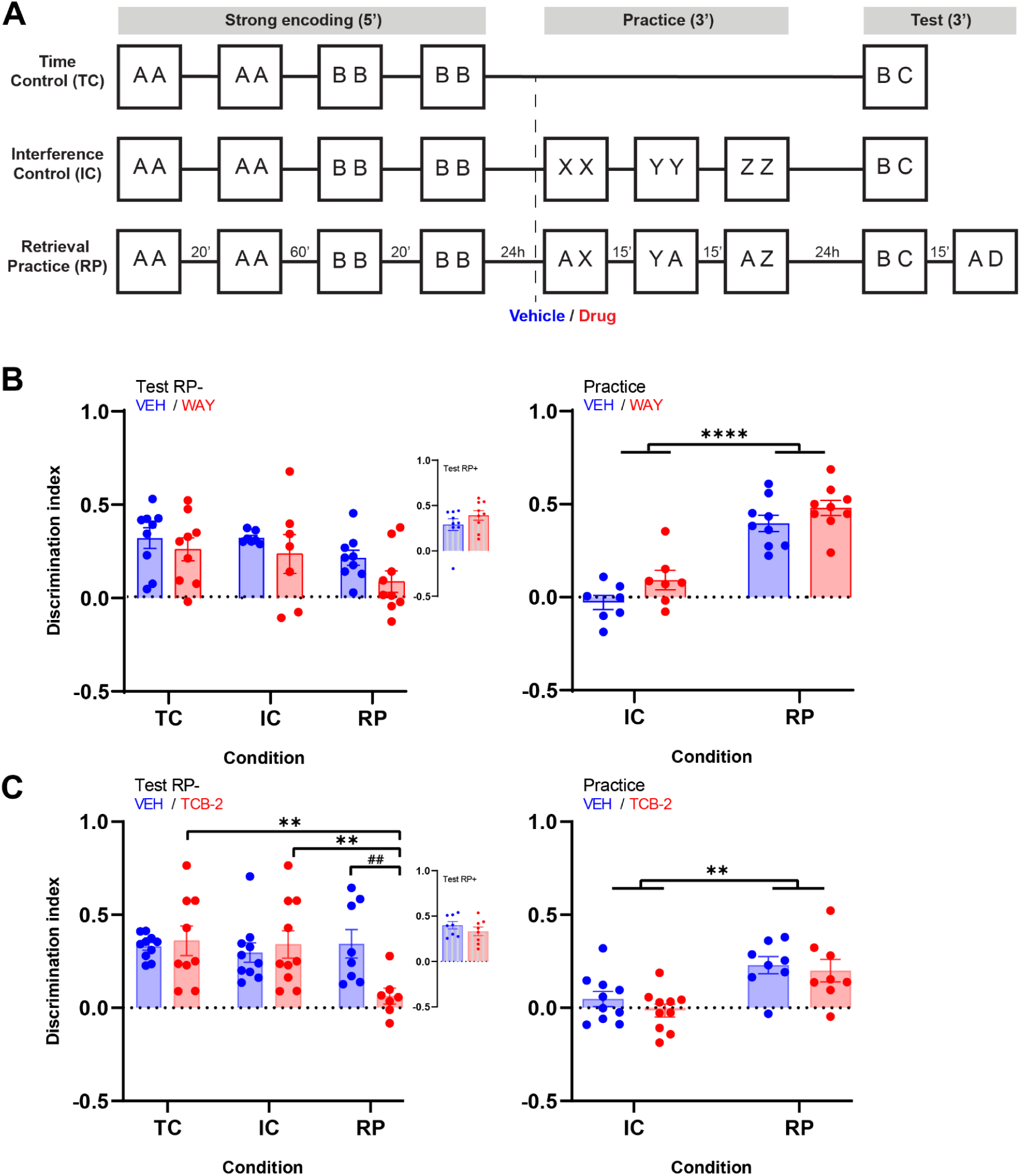
Serotonergic manipulations under conditions that do not normally induce RIF. **A.** Schematic of the behavioral protocol. In contrast to Figure 1, this protocol doubled the number of encoding sessions and separated encoding from practice by 24 hours. **B.** Left: Discrimination index ± SEM at test for the 5-HT1AR antagonist experiment (WAY; n = 25); the Condition_RP_ x Drug_WAY_ interaction was not significant. Inset: Discrimination index for the practiced object. Right: Discrimination index during practice for IC and RP groups. **C.** Left: Discrimination index at test for the 5-HT2AR agonist experiment (TCB-2; n = 28). Asterisks (*) indicate differences between conditions within each treatment and number signs (#) indicate differences between treatments within conditions (Bonferroni post hoc). Inset: Discrimination index for the practiced object. Right: Discrimination index during practice for IC and RP groups. *p < 0.05, **p < 0.01, ***p < 0.001, ****p < 0.0001.

Because activation of mPFC 5-HT1AR impaired RIF in the standard protocol, we first tested whether blocking this inhibitory receptor could facilitate forgetting. Animals received intra-mPFC infusions of vehicle or the selective 5-HT1AR antagonist WAY 100135 (WAY) before the practice phase. In the final test, we found no significant Condition_RP_ x Drug_WAY_ interaction (β = −0.07, _95%_CI [−0.29, 0.16], t_(22)_ = −0.58, p = .57; Condition_IC_ × Drug_WAY_: β = −0.03, _95%_CI [−0.27, 0.22], t_(22)_ = −0.21, p = .84), indicating that 5-HT1AR blockade was not sufficient to induce RIF in this modified protocol. Likewise, we found no main effect of Drug (Drug_WAY_: β = −0.06, _95%_CI [−0.22, 0.11], t_(22)_ = −0.74, p = .47). No treatment differences were observed for memory of the practiced object (A, RP+; Drug_WAY_: β = 0.10, _95%_CI [−0.03, 0.24], t_(8)_ = 1.75, p = .12; Fig. 2B, left, inset).

We also analyzed performance during the practice phase to determine whether retrieval practice was affected by the treatment. As expected from the structure of the task, RP animals showed higher DI values than IC animals (Condition_RP_: β = 0.43, _95%_CI [0.29, 0.56], t_(14)_ = 6.80, p < .001; [IC−RP]: mean = -0.407, SE = 0.044, t_(14)_ = -9.188, p < .0001), regardless of the pharmacological treatment, indicating that WAY did not alter performance during retrieval practice (Fig. 2B, right).

We next asked whether the role of the 5-HT2AR in RIF extends beyond necessity to sufficiency. Because blocking mPFC 5-HT2AR impaired RIF for the competitor object, we tested whether pharmacological activation of this receptor could be sufficient to promote forgetting. Under these conditions, animals received intra-mPFC infusions of vehicle or the selective 5-HT2AR agonist TCB-2 before the practice phase.

If activation of the 5-HT2AR is sufficient to promote forgetting of a competing memory, only RP animals treated with TCB-2 should fail to discriminate between the competitor object (B, RP−) and the novel object in the final test. As predicted, RP animals infused with TCB-2 explored both objects similarly, unlike the IC and TC groups ([TC−RP]_TCB-2_: mean = 0.291, SE = 0.079, t_(25)_ = 3.683, p < .01; [IC−RP]_TCB-2_: mean = 0.282, SE = 0.079, t_(25)_ = 3.572, p < .01). Consistent with this pattern, we found a significant Condition_RP_ x Drug_TCB-2_ interaction (β = −0.31, _95%_CI [−0.53, −0.08], t_(25)_ = −2.80, p = .01). Post hoc comparisons showed that TCB-2 reduced DI only in the RP condition (RP_VEH−TCB-2_: mean = 0.286, SE = 0.081, t_(25)_ = 3.520, p < .01), with no differences in the IC (IC_VEH−TCB-2_: mean = −0.043, SE = 0.073, t_(25)_ = −0.596, p > .99) or TC (TC_VEH−TCB-2_: mean = −0.019, SE = 0.073, t_(25)_ = −0.258, p > .99) conditions (Fig. 2B, left). No treatment differences were observed in memory for the practiced object (A, RP+) (Fig. 2B, left, inset).

We also analyzed performance during the practice phase to determine whether the test effect could be explained by differences during retrieval practice. As in the previous experiments, TCB-2 did not alter performance during this phase. We detected only a significant main effect of condition (β = 0.18, _95%_CI [0.05, 0.32], t_(16)_ = 2.83, p < .05; [IC−RP]: mean = -0.198, SE = 0.05, t_(16)_ = -3.936, p < .01), indicating that RP animals discriminated more strongly than IC animals, as expected from the structure of the task. Thus, infusion of TCB-2 before retrieval practice did not appear to affect attention, exploratory drive, or motivation (Fig. 2B, right).

### PI3K mediates the 5-HT2AR effect on RIF in the CPFm during Retrieval Practice

We next examined whether the PI3K/AKT pathway contributes to the control of RIF. Although PI3K/AKT is not considered the canonical signaling pathway of the 5-HT2AR, it plays an important role in neuronal homeostasis, plasticity, and long-term memory, making it a plausible candidate mechanism in this process (Horwood et al., 2006). To test this possibility, we used the same behavioral design as in the 5-HT2AR blockade experiment. Animals were trained in the RIF task, randomly assigned to the RP, IC, or TC conditions, and received intra-mPFC infusions of the PI3K inhibitor LY294002 (LY) or vehicle before the practice phase (Fig. 3A).

**Figure 3.**
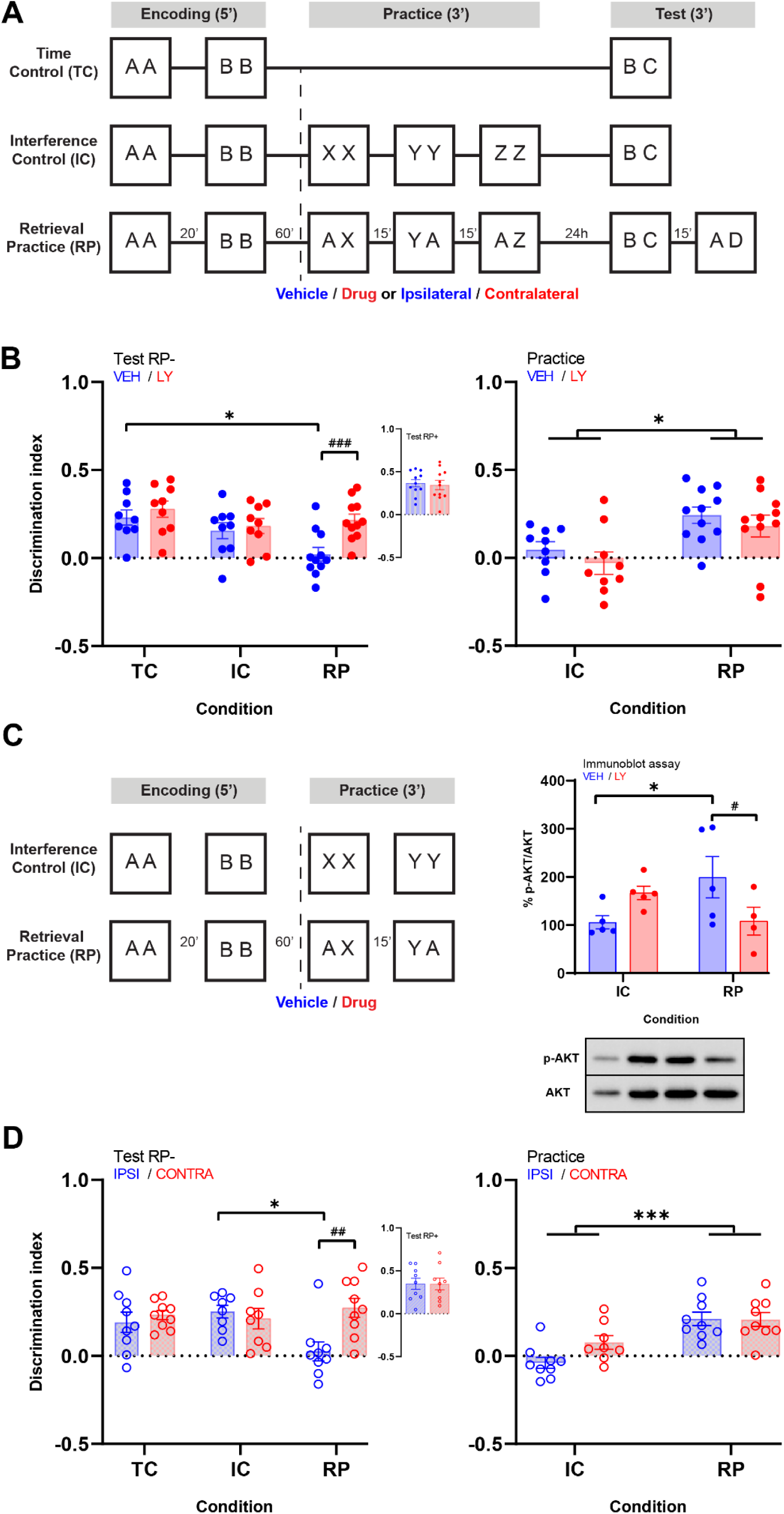
PI3K is involved in 5-HT2AR signaling during retrieval practice. **A.** Schematic of the behavioral protocol. VEH or LY294002 (LY) was infused into the mPFC 15 min before practice. In the disconnection experiment, LY and MDL were infused ipsilaterally (IPSI) or contralaterally (CONTRA). **B.** Left: Discrimination index ± SEM at test in the LY experiment (n = 29). Inset: Discrimination index for the practiced object. Right: Discrimination index during practice for IC and RP groups. **C.** Left: Schematic of the abbreviated protocol for the immunoblot assay; VEH or LY was infused before practice and tissue was collected after the second session. Right: p-AKT/AKT levels ± SEM in IC and RP groups (n = 19) with a representative immunoblot; asterisks indicate planned comparisons. **D.** Left: Discrimination index at test in the disconnection experiment (n = 26). Inset: Discrimination index for the practiced object. Right: Discrimination index during practice for IC and RP groups. Asterisks (*) indicate significant differences between conditions within each treatment and number signs (#) indicate significant differences between treatments within conditions (Bonferroni post hoc). *p < 0.05, **p < 0.01, ***p < 0.001, ****p < 0.0001.

If PI3K activity is recruited during selective retrieval to promote forgetting of competing memories, LY infusion should impair RIF. The results supported this prediction (Fig. 3B, left). We found a significant Condition_RP_ x Drug_LY_ interaction (β = 0.13, _95%_CI [0.01, 0.25], t_(25)_ = 2.18, p < .05), and post hoc comparisons showed that LY increased DI selectively in the RP condition (RP_VEH−LY_: mean = −0.193, SE = 0.039, t_(25)_ = −4.907, p < .001). No drug effect was observed in the control conditions (IC_VEH−LY_: mean = −0.026, SE = 0.044, t_(25)_ = −0.602, p > .99; TC_VEH−LY_: mean = −0.063, SE = 0.045, t_(25)_ = −1.408, p > .5). Consistent with RIF, VEH-treated animals showed a lower DI in the RP condition than in the TC condition ([TC−RP]_VEH_: mean = 0.202, SE = 0.058, t_(25)_ = 3.484, p < .05). Thus, PI3K inhibition impaired RIF specifically, rather than producing a general long-term memory deficit. No treatment differences were observed for memory of the practiced object (Fig. 3B, left, inset).

We also analyzed behavior during the practice phase to determine whether the test-phase effect could be explained by altered performance during retrieval practice. As in the previous experiments, LY infusion did not affect discrimination during this phase (Fig. 3B, right); we detected only a significant effect of condition (β = 0.20, _95%_CI [0.03, 0.36], t_(18)_ = 2.50, p < .05; [IC−RP]: mean = -0.204, SE = 0.056, t_(18)_ = -3.676, p < .01).

To explore whether the behavioral effects observed during retrieval practice were associated with changes in PI3K/AKT signaling, we quantified phosphorylated AKT levels by Western blot in IC and RP animals treated with either vehicle or LY. If retrieval practice recruits PI3K/AKT signaling, phosphorylated AKT levels should increase in RP animals under vehicle treatment, and LY administration should prevent this increase. The results supported this prediction (Fig. 3C). We found a significant Condition x Drug interaction (F_(1,15)_ = 7.46, p < .05). Planned contrasts showed that under vehicle treatment, phosphorylated AKT levels were higher in RP animals than in IC controls ([IC−RP]_VEH_: mean = −93.65, SE = 38.28, t_(15)_ = −2.45, p < .05). LY reduced phosphorylated AKT levels in the RP condition ([VEH−LY]_RP_: mean = 91.32, SE = 40.60, t_(15)_ = 2.25, p < .05). No effect of LY was observed in the IC condition ([VEH−LY]_IC_: mean = −61.05, SE = 38.28, t_(15)_ = −1.60, p = .132), and no difference between IC and RP was detected under LY treatment ([IC−RP]_LY_: mean = 58.72, SE = 40.60, t_(15)_ = 1.45, p = .169). Thus, retrieval practice increased PI3K/AKT signaling in the mPFC, and this effect was prevented by PI3K inhibition. LY also did not alter performance during the practice phase in the abbreviated protocol used for biochemical analyses. We detected only a significant effect of condition (Condition_RP_: β = 0.26, _95%_CI [0.05, 0.48], t_(15)_ = 2.68, p < .05; [IC−RP]: mean = -0.331, SE = 0.073, t_(15)_ = -4.513, p < .0001), indicating that LY administration did not interfere with the expression of the practiced memory or with exploratory behavior during this phase

We then asked whether PI3K mediates 5-HT2AR modulation of RIF using a molecular disconnection approach. This strategy is based on the logic of classic disconnection experiments, which assess functional interactions by disrupting two elements within the same circuit or process in opposite hemispheres (Jo and Lee, 2010; Bekinschtein et al., 2013; Miranda et al., 2017, 2018). If 5-HT2AR signaling and PI3K activity interact within the mPFC to support RIF, then ipsilateral infusions should preserve function in the intact hemisphere, whereas contralateral infusions should disrupt the behavioral effect. To test this, cannulated rats were trained in the RIF task (Fig. 3A) and, before the practice phase, received intra-mPFC infusions of MDL and LY either in the same hemisphere (ipsilateral) or in opposite hemispheres (contralateral). When memory for the competitor object (B, RP−) was tested, animals receiving contralateral infusions could distinguish between the novel object and the RP− object, indicating disruption of RIF (Fig. 3D, left). Consistent with this result, we found a significant Condition_RP_ x Drug_CONTRA_ interaction (β = 0.21, _95%_CI [0.02, 0.40], t_(22)_ = 2.25, p < .05). Post hoc comparisons showed that contralateral infusions increased DI selectively in the RP condition (RP_IPSI−CONTRA_: mean = −0.247, SE = 0.065, t_(23)_ = −3.799, p < .01), with no effect in the IC (IC_IPSI−CONTRA_: mean = −0.040, SE = 0.069, t_(23)_ = 0.577, p > .99) or TC (TC_IPSI−CONTRA_: mean = −0.040, SE = 0.065, t_(23)_ = −0.621, p > .99) conditions. Consistent with RIF, ipsilaterally infused animals showed a lower DI in the RP condition than in the IC condition ([IC−RP]_IPSI_: mean = 0.225, SE = 0.070, t_(23)_ = 3.205, p < .05). No differences were observed between infusion schemes for memory of the practiced object (Fig. 3D, left, inset). These findings support the hypothesis that PI3K activity mediates, at least in part, 5-HT2AR modulation of RIF.

Finally, neither infusion scheme affected performance during the practice phase (Fig. 3D, right). We detected only a significant effect of condition (Condition_RP_: β = 0.25, _95%_CI [0.13, 0.36], t_(15)_ = 4.54, p < .001; [IC−RP]: mean = -0.188, SE = 0.038, t_(15)_ = -4.912, p < .001), indicating that the manipulations did not interfere with expression of the practiced memory or with the ability to explore novel objects.

### Src contributes to RIF during retrieval practice, but is not functionally linked to 5-HT2AR signaling

We next examined whether Src signaling contributes to RIF. Src family kinases (here referred to as Src to denote the SFK family) are non-receptor tyrosine kinases widely expressed in the brain and involved in multiple neuronal processes (Omri et al., 1996; Kalia et al., 2004; Bongiorno-Borbone et al., 2005). In particular, Src and Fyn have been implicated in the modulation of synaptic transmission and plasticity (Kalia et al., 2004; Ohnishi et al., 2011; Schenone et al., 2011). In addition, it has been shown that serotonin promotes formation of a 5-HT2AR signaling complex involving β-arrestin2, PI3K, Src, and AKT, and that blocking any of these components, including PI3K, prevents full expression of the head-twitch response (Schmid and Bohn, 2010). Given the role of Src signaling in neuronal plasticity and the effects of PI3K inhibition on RIF, we asked whether Src inhibition would similarly disrupt RIF.

To test this, animals were trained in the RIF task and received intra-mPFC infusions of the Src inhibitor PP2 or vehicle before the practice phase (Fig. 4A). Consistent with RIF, VEH-treated animals showed a lower DI in the RP condition than in the IC and TC conditions ([TC−RP]_VEH_: mean = 0.337, SE = 0.051, t_(17)_ = 6.559, p < .0001; [IC−RP]_VEH_: mean = 0.340, SE = 0.053, t_(17)_ = 6.362, p < .0001). Supporting our hypothesis, we found a significant Condition_RP_ x Drug_PP2_ interaction (β = 0.19, _95%_CI [0.04, 0.34], t_(17)_ = 2.60, p < .05), and post hoc comparisons trend towards inhibiting RIF expression. (RP_VEH−PP2_: mean = −0.127, SE = 0.051, t_(17)_ = −2.477, *p = .07*). No significant effect in the IC (IC_VEH−PP2_: mean = 0.094, SE = 0.055, t_(17)_ = 1.688, p > .5) or TC (TC_VEH−PP2_: mean = 0.062, SE = 0.051, t_(17)_ = 1.205, p > .5) conditions. These results point to a possible role of Src in RIF without producing a general long-term memory deficit (Fig. 4B, left). No treatment-related differences were observed for memory of the practiced object (Fig. 4B, left, inset).

**Figure 4.**
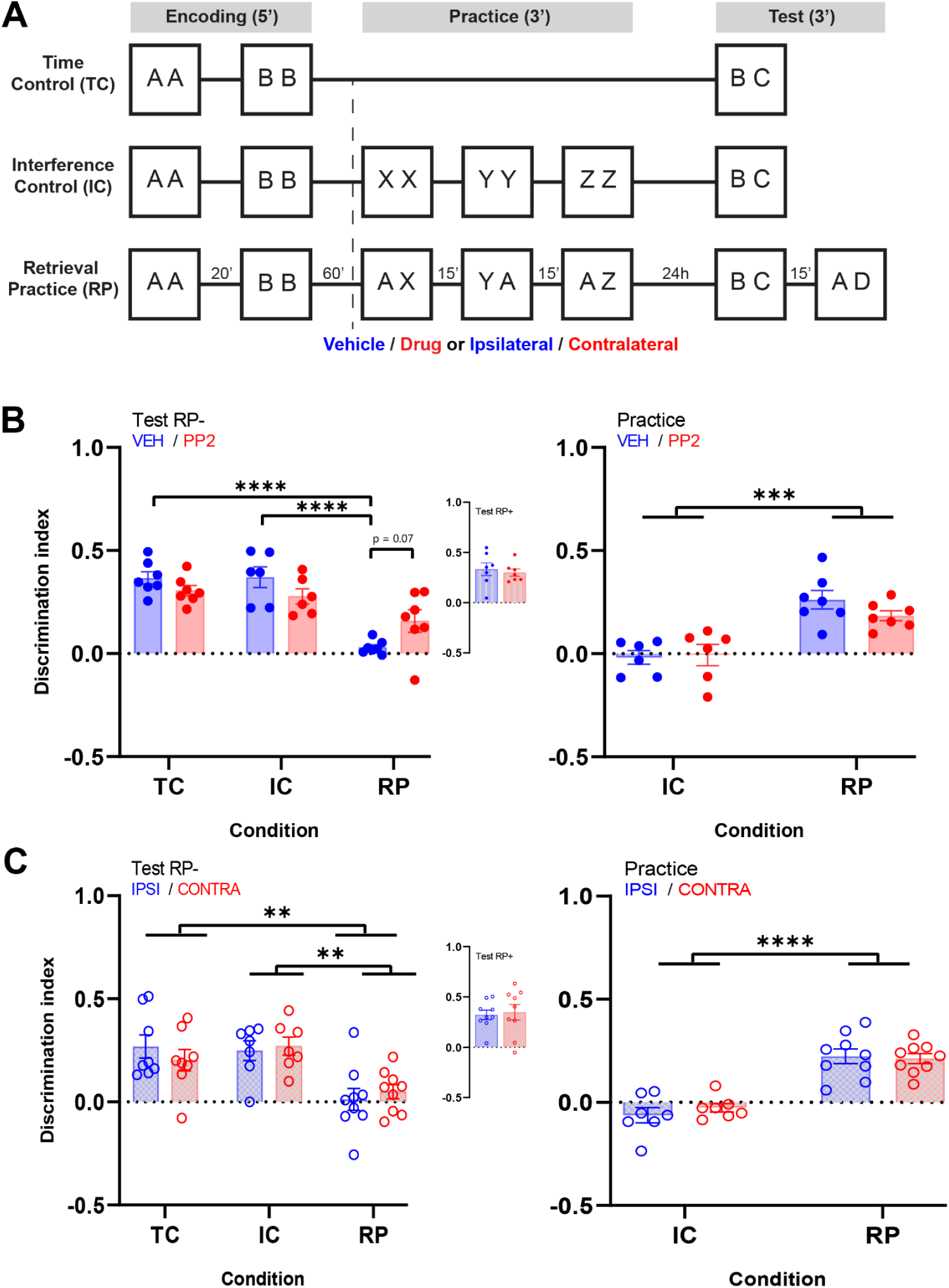
Rol of Src activity during retrieval practice for RIF. **A.** Schematic of the behavioral protocol. VEH or the Src inhibitor PP2 was infused into the mPFC 15 min before practice. In the disconnection experiment, PP2 and MDL were infused ipsilaterally (IPSI) or contralaterally (CONTRA). **B.** Left: Discrimination index ± SEM at test in the PP2 experiment (n = 20). Inset: Discrimination index for the practiced object. Right: Discrimination index during practice for IC and RP groups. **C.** Left: Discrimination index at test in the disconnection experiment (n = 24); asterisks indicate the main effect of condition, as the interaction was not significant. Inset: Discrimination index for the practiced object. Right: Discrimination index during practice for IC and RP groups. *p < 0.05, **p < 0.01, ***p < 0.001, ****p < 0.0001.

We also analyzed behavior during the practice phase As in the previous experiments, PP2 did not affect performance during this phase; we detected only a significant effect of condition (β = 0.28, _95%_CI [0.16, 0.40], t_(11)_ = 5.03, p < .001; [IC−RP]: mean = -0.235, SE = 0.039, t_(11)_ = -5.969, p < .001), indicating the expected difference between RP and IC animals (Fig. 4B, right). Analysis of total exploration time revealed a significant main effect of Drug (β = −22.77, _95%_CI [−41.59, −3.95], t_(11)_ = −2.66, p < .05; [VEH-PP2]: mean = 31.555, SE = 5.826, t_(11)_ = 5.416, p < .001), with PP2-treated animals exploring less overall during the practice phase. However, since this effect was not specific to the RP condition (Condition_RP_ × Drug_PP2_ interaction: p = .160), it cannot account for the selective impairment of RIF observed at test.

We then tested the functional interaction between Src and the 5-HT2AR using a molecular disconnection approach similar to that described above. Cannulated rats were trained in the RIF paradigm (Fig. 4A) and, before the practice phase, received intra-mPFC infusions of MDL and PP2 either in the same hemisphere (IPSI) or in opposite hemispheres (CONTRA). Contrary to our prediction, we found no significant Condition_RP_ × Drug_CONTRA_ interaction in the competitor object memory test. Instead, we observed only a main effect of condition, indicating preserved RIF regardless of infusion scheme (Condition_RP_: β = −0.26, _95%_CI [−0.40, −0.12], t_(21)_ = −3.84, p < .001; [IC−RP]: mean = 0.229, SE = 0.061, t_(21)_ = 3.754, p < .01; [TC−RP]: mean = 0.205, SE = 0.059, t_(21)_ = 3.490, p < .001). Thus, we found no evidence that Src signaling functionally interacts with the 5-HT2A receptor during retrieval practice to support RIF (Fig. 4C, left). No differences between infusion schemes were observed for memory of the practiced object (Fig. 4C, left, inset).

Finally, the pharmacological manipulations did not alter behavior during the practice phase of the disconnection experiment. We detected only a significant effect of condition (β = 0.29, _95%_CI [0.19, 0.38], t_(14)_ = 6.55, p < .001; [IC−RP]: mean = -0.262, SE = 0.036, t_(14)_ = -7.332, p < .0001), again indicating the expected difference between RP and IC animals (Fig. 4C, right).

## Discussion

### Active forgetting and serotonergic modulation of retrieval-induced forgetting

Historically, neurobiological research on memory has focused on understanding how memories are encoded, stored, and retrieved, whereas forgetting has often been regarded as memory failure (Davis and Zhong, 2017; Medina, 2018). However, emerging perspectives suggest that distinct mechanisms may promote forgetting to achieve a range of functional ends (Davis and Zhong, 2017; Richards and Frankland, 2017; Anderson and Hulbert, 2021; Ryan and Frankland, 2022). RIF is among active processes that promote forgetting (Anderson and Hulbert, 2021). Cognitive models propose that RIF relies on inhibitory control processes, wherein attempts to retrieve target memories inadvertently activate competing, contextually irrelevant memories, which are then suppressed, reducing interference and facilitating efficient retrieval. In addition to facilitating access to the desired memory, the persisting effects of this mechanism would enhance future retrievals of the target memory, preparing the organism for similar circumstances and optimizing behavioral responses. In this framework, forgetting can be conceptualized as an active mechanism that improves behavioral flexibility and mechanistically relies on cortical inhibitory processes.

Traditionally studied in the context of cognitive control and mood disorders (Clarke et al., 2007; Robbins, 2016; Luo et al., 2024; Nutt et al., 2024), the serotonergic system has more recently emerged as a key modulator of learning and memory mechanisms, particularly within the mPFC (Pacheco et al., 2012; Zhang and Stackman, 2015; Tortora et al., 2023). Here, we extend the role of the serotonergic system to include its involvement in the adaptive forgetting of competing memories. Our findings indicate that serotonergic modulation in the mPFC is required during repeated retrieval attempts, suggesting that serotonin participates directly in the inhibitory control processes that underlie RIF.

Pharmacological manipulations revealed a receptor-specific contribution to this effect. Blocking the 5-HT2AR in the mPFC during the practice phase blocked RIF without altering memory for the practiced object (Fig. 1C, *center*), supporting a necessary role for 5-HT2AR activity in this process. Conversely, activating the 5-HT2AR in a protocol that does not normally induce RIF was sufficient to elicit forgetting of the competing memory.

Our findings also indicate that the 5-HT1AR modulates this phenomenon, although in a different manner. Though activation of 5-HT1A impaired forgetting, inhibition has no effect (Fig. 1C, *left*). Thus, unlike the 5-HT2AR, the 5-HT1AR does not appear to drive RIF directly, but rather to constrain or modulate the cortical dynamics that permit it.

5-HT2CR are also expressed in the mPFC, and it has been demonstrated that MDL11,939 exhibits some affinity for them (Pehek et al., 2006). However, infusion of SB242084, a potent and selective 5-HT2CR antagonist (Kennett et al., 1997), did not affect performance in the RIF task. Together, these findings suggest that serotonergic modulation of RIF depends primarily on the 5-HT2AR, is disrupted by 5-HT1AR activation, and does not detectably involve the 5-HT2CR under our experimental conditions.

### A serotonergic control mechanism in the mPFC

The prominent serotonergic innervation of the mPFC, together with the heterogeneous distribution of serotonin receptors, provides a framework for understanding these effects. Both 5-HT1A and 5-HT2A receptors are highly expressed in this region, where they exert opposing influences on neuronal excitability: 5-HT1AR activation typically reduces firing, whereas 5-HT2AR activation enhances it. This push-pull dynamic is thought to regulate prefrontal network activity and cognitive processing. In this context, our results suggest that RIF emerges from a balance in which 5-HT2AR activation promotes the suppression of competing representations, while 5-HT1AR activity limits excessive excitation, thereby stabilizing network function. Such coordinated modulation may be critical for implementing selective inhibitory control without disrupting overall circuit integrity (Araneda and Andrade, 1991; Amargós-Bosch et al., 2004; Andrade, 2011; Mengod et al., 2015). At the behavioral level, 5-HT1A receptors have been implicated in several cognitive domains, including attention, executive function, working memory, fear processing, and recognition memory (Carli et al., 1992; Winstanley et al., 2003; Stiedl et al., 2015; Nikolaus et al., 2024). Importantly, pharmacological studies indicate that 5-HT1AR activation often impairs cognitive performance, particularly in tasks requiring memory retrieval (Pitsikas et al., 2005), whereas blockade of these receptors produces more variable effects, ranging from facilitation to no detectable change depending on task demands and baseline network state (Meneses and Perez-Garcia, 2007). For example, 5-HT1AR activation has been reported to disrupt recognition memory and working memory performance, while antagonism does not consistently enhance these functions (Meneses and Perez-Garcia, 2007).

Notably, 5-HT2A receptors are implicated in cognitive functions that require the control of prepotent responses (Boulougouris et al., 2008; Robinson et al., 2008). Our results extend this role to the control of memory retrieval, showing that serotonergic modulation of RIF operates through the coordinated activation of specific receptor subtypes, 5-HT1A and 5-HT2A, but not 5-HT2C. Specifically, blocking mPFC 5-HT2AR prior to the retrieval practice phase impaired RIF, whereas activating these receptors immediately before retrieval practice enhanced it.

What are the mechanisms by which serotonin modulates RIF? Active inhibition of downstream areas like the medial temporal lobe might be triggered by the mPFC inhibiting the memory of the competitor object (Anderson et al., 2016) in a serotonin-dependent manner. Temporary interactions between low- and high-frequency brain waves are proposed to coordinate network activity across the cortical surface during cognitive tasks (Canolty et al., 2006). In the PFC, 5-HT2A and 5-HT1A receptors jointly contribute to generating gamma oscillations (Puig and Gulledge, 2011), while exerting opposing influences on gamma wave power: 5-HT1AR antagonism enhances cortical gamma activity, whereas 5-HT2AR antagonism disrupts gamma wave synchronization (Celada et al., 2004). Comparable effects on gamma and theta rhythms have been reported in mice, suggesting that blocking 5-HT2A or activating 5-HT1A receptors could disrupt the balance essential for serotonergic regulation of cortical network dynamics during active inhibition. Human studies further show that reduced theta and gamma power are linked to the forgetting of unpracticed items (Spitzer et al., 2009), suggesting that serotonin may enhance the expression of these rhythms and thereby contribute to the inhibition of unpracticed objects.

### Retrieval-induced forgetting and other forms of retrieval-based inhibition

Interestingly, previous work using the Object-in-Context task has shown that during retrieval, when multiple memory representations compete, the less relevant memory can be inhibited. This suppression resembles the inhibitory processes engaged during the initial practice sessions of the RIF paradigm, in which selective retrieval of certain items leads to competition among related memories. However, there seems to be a key divergence between these paradigms in terms of the persistence of forgetting. In the Object-in-Context task, when memory for the originally encoded associations is assessed 24 hours later, those memories appear to remain intact, suggesting that the inhibition observed during retrieval is temporary and reversible. In contrast, in the RIF paradigm, memory for the competing item (e.g., object B) is typically no longer accessible after 24 hours, indicating a more lasting form of inhibition. This contrast raises important questions regarding the nature and mechanisms of inhibitory control in memory. Are the forms of inhibition engaged in these two paradigms fundamentally different in their temporal dynamics? Might they reflect distinct neural processes, even if both involve activation of the 5-HT2AR in the mPFC? While our data suggest a shared reliance on this receptor, the differences in the persistence of memory suppression point to potentially divergent downstream mechanisms. Future studies will be necessary to determine whether these two phenomena represent qualitatively different forms of inhibition or different outcomes of a shared mechanism, modulated by contextual factors.

### PI3K/AKT signaling as a downstream pathway of 5-HT2A receptor modulation

5-HT2AR is a GPCR with a complex system of signaling pathways. The classical pathway is G_q/11_-mediated activation of PLC and Ca^2+^ release from the endoplasmic reticulum. However, studies in heterologous expression systems identified several other signaling transduction pathways coupled to the receptor. Different pathways appear to be selectively activated depending on the type of agonist used to test them (Berg et al., 1998; Porter et al., 1999; Kurrasch-Orbaugh et al., 2003). This phenomenon, which seems to be a feature of GPCRs, is known as biased agonism. In recent years, efforts to identify the signaling pathway downstream of 5-HT2AR have received considerable attention as a renewed interest in hallucinogenic drugs as therapeutic agents has increased. However, the endogenous 5-HT2AR ligand, serotonin, can also activate different signaling pathways depending on its release kinetics and the cellular milieu. Activation of 5-HT2AR can induce the ERK and tyrosine kinase pathways, as well as the β-arrestin2/PI3K/Src/AKT cascade (Schmid and Bohn, 2010). Because different pathways may contribute uniquely to various cognitive processes, we focused on the PI3K/AKT pathway as a potential mechanism involved in the modulation of RIF by 5-HT2AR. The PI3K/AKT pathway is a critical regulator of brain function, playing essential roles in neuronal homeostasis, growth, and plasticity. Although the PI3K/AKT pathway is not the canonical 5-HT2AR signaling route, its involvement is particularly intriguing due to the relationship between AKT and long-term memory processes (Horwood et al., 2006).

In this work, we found that PI3K activity in the mPFC is necessary for RIF expression, with AKT as a downstream mediator: AKT phosphorylation changed between treatments, indicating that this pathway is recruited during the retrieval practice phase. We also found a functional interaction between 5-HT2AR and PI3K, which is essential for 5-HT2AR activation during the retrieval practice phase. Independent of the complete pathway involved, our results support PI3K as a molecular bridge between 5-HT2AR signaling and active forgetting.

### Src contributes to RIF but is not directly linked to 5-HT2AR signaling

The 5-HT2AR has been shown to couple to activation of Src family kinases in neuronal cells (González-Maeso et al., 2007; Schmid and Bohn, 2010; Tang and Trussell, 2015). Accumulating evidence suggests that Src kinase plays a role in learning and memory, potentially through phosphorylation of NMDA receptor subunits and modulation of synaptic plasticity. However, its contribution remains complex and not fully understood. For instance, increased Src activity has been linked to enhanced hippocampal long-term potentiation and improved spatial memory (Wang et al., 2018), and spatial learning has been associated with upregulation of Src mRNA in area CA3 (Zhao et al., 2000). In contrast, Src overexpression can disrupt excitatory synaptic transmission in CA3 and impair fear memory (Yan et al., 2017), highlighting that its effects may depend on the level and context of activation. Even less is known about its potential involvement in active forgetting mechanisms, and virtually nothing about its role in retrieval-induced forgetting (RIF).

Because Src participates in a shared pathway with 5-HT2AR and PI3K, we hypothesized that it might also contribute to RIF. Our results show that inhibiting Src during the practice phase interferes with RIF expression, indicating that Src activity contributes to this process. However, Src does not appear to act downstream of 5-HT2AR during retrieval, pointing instead to a parallel signaling pathway. Because Src family kinases can be recruited by multiple upstream signals, it is conceivable that an unidentified upstream regulator activates Src independently of 5-HT2AR, ultimately converging on AKT. This possibility remains to be addressed in future studies.

Few studies have explored the direct interaction between GPCRs and Src or their activation mechanisms. Like arrestins, Src offers an alternative signaling pathway for GPCRs, contributing to their physiological functions. Further research is needed to clarify the specific pathways mediated by Src and their role in GPCR signaling (Berndt and Liebscher, 2021).

### Interactions with dopamine and broader neuromodulatory control

Previously, we have shown that dopamine acting on D1Rs in the mPFC modulates the control processes involved in RIF in rats (Gallo et al., 2022). The serotonergic and dopaminergic systems often interact, influencing various functions including cognition and mood regulation (De Deurwaerdère et al., 2018, 2021). Although dopamine and serotonin have similar functional relevance, the mechanisms by which serotonin affects behavior are less well understood than those of dopamine (De Deurwaerdère et al., 2021). Regarding the serotonin 5-HT2AR, it has been proposed that it may influence dopamine release. Systemic administration or local infusion of DOI, a selective 5-HT2AR agonist, in the PFC results in an increase in local dopamine, and this effect can be blocked by selective 5-HT2AR antagonists (Pehek et al., 2001, 2006; Bortolozzi et al., 2005). In addition, 5-HT2AR is expressed in approximately 55% of pyramidal neurons projecting to the ventral tegmental area (Vázquez-Borsetti et al., 2009), and half of these neurons also synapse in the dorsal raphe nucleus (DR), suggesting that mPFC 5-HT2A receptors might be involved in the concerted modulation of ascending serotonergic and dopaminergic activity (Vázquez-Borsetti et al., 2011). Dopamine and serotonin might work in concert during RIF, but further studies are necessary to test this hypothesis.

### Clinical implications for depression and intrusive cognition

Individuals with Major Depressive Disorder (MDD) exhibit specific characteristics in episodic memory retrieval and the emotional valence of recalled events, which may stem from difficulties in inhibiting irrelevant information (Lemogne et al., 2006). Notably, both anxiety and depressive disorders are associated with reduced RIF (Groome and Sterkaj, 2010), suggesting that the inhibitory processes crucial for RIF are disrupted in these conditions. RIF is known to depend on PFC activity (Kuhl et al., 2007; Wimber et al., 2015), and the functionality of this region is markedly altered in depression (Downar and Daskalakis, 2013; Riva-Posse et al., 2018). Could the impaired inhibitory control observed in individuals with MDD (Giebl et al., 2016) be the mechanism responsible for their diminished RIF? Our findings further position serotonin 5-HT2A and 5-HT1A receptors, both implicated in depressive phenotypes (Celada et al., 2004; Savitz et al., 2009; Albert and François, 2010), as critical contributors to the processes underlying RIF. Examining how antidepressant treatment affects RIF performance could clarify the interactions between depressive states and memory processes.

Beyond depression, deficits in serotonergic modulation and inhibitory control have been implicated in anxiety, obsessive-compulsive disorder, and impulsivity-related conditions. If serotonergic activity contributes to active forgetting via inhibitory control, as the present findings indicate, deficits in serotonin may partially underlie the inability to down-regulate intrusive thoughts in pathological worry, rumination, and obsessive thinking. This is consistent with evidence that the success of thought suppression rests on active forgetting (Mamat and Anderson, 2023; Anderson et al., 2025). More broadly, by positioning the serotonergic system in the mPFC as a critical regulator of memory suppression, these findings contribute to a general framework for understanding how active forgetting supports cognitive flexibility and adaptive behavior in dynamic environments.

Taken together, our findings establish a causal role for serotonergic signaling in the mPFC in retrieval-induced forgetting, identifying 5-HT2AR activation as necessary and sufficient for the inhibitory control mechanism recruited during retrieval practice. The pharmacological effects were strictly condition-specific: no changes were detected when the same compounds were administered before exposure to novel objects (IC) or during the equivalent time interval in the home cage (TC), ruling out contributions of novelty, salience, or nonspecific effects on exploratory behavior. This selectivity constrains the interpretation considerably: 5-HT2AR signaling does not modulate memory storage or retrieval *per se*, but rather the suppression mechanism engaged specifically when a competing memory trace is reactivated. The disconnection experiments further implicate a functionally integrated intrahemispheric circuit as the substrate of this modulation, positioning the mPFC not merely as a site of serotonergic action but as an active node in a circuit that executes retrieval-induced suppression. Finally, the PI3K/AKT pathway emerges as a downstream effector of 5-HT2AR signaling during retrieval practice, providing a first molecular substrate for the inhibitory control mechanism underlying RIF.

## Materials and Methods

### Ethics statement

All experimental procedures were performed according to institutional regulations and following those of the National Animal Care and Use Committee of the University of Buenos Aires and Favaloro University (CICUAL Presentation DCT0204-16 / PICT 2015-0110). All efforts were made to minimize the number of animals used and their suffering.

### Subjects

We used 268 (116 male and 152 female) Wistar rats from our breeding colony, weighing 200-300 grams at the beginning of each experiment. The animals were kept in groups of 4 to 5 individuals with access to food and water ad libitum, under a 12-hour light/dark cycle (7 a.m.–7 p.m.) and a temperature between 20 and 23 °C. The number of animals for each experiment is reported in the corresponding results section. All experiments were performed during the light phase of the cycle. In all cases, the animals were handled for two consecutive days before the start of the experiment.

### Retrieval-Induced Forgetting task

#### Overview

We used the task developed by Bekinschtein et al. (Bekinschtein et al., 2018), which adapts the spontaneous object recognition procedure to include phases analogous to those used in human RIF studies (Anderson et al., 1994; Ciranni and Shimamura, 1999; Wimber et al., 2015). The task comprises three experimental conditions, Retrieval Practice (RP), Interference Control (IC), and Time Control (TC), each consisting of three phases (Fig. 1A). The RP condition is designed to induce the forgetting of an object memory triggered by the successive retrieval of a related object memory. The IC condition allows for assessing the effect of repeated presentation of objects on the recall of the studied objects. The TC condition evaluates the possible effect that the time elapsed from learning to evaluation could have on the retention of the memory of the objects.

In all experiments, animals were randomly assigned to one of the three experimental conditions following the shaping phase (see below). The order of exposure to each treatment (drug/vehicle or practice length) was pseudorandomized, and experiments were conducted over a two-week period.

#### The general retrieval practice paradigm

Shaping: To familiarize the animals with the general features of the task, they were first placed in an arena for 10 minutes and then exposed to a pair of objects in the same arena for 3 minutes. Neither the arena nor the objects used during shaping were employed in the actual task, which began 24 to 48 hours later.

Day 1 - Context Habituation: Animals were allowed to freely explore a novel context for 10 minutes.

Day 2:

- Encoding: Twenty-four hours after context habituation, animals were reintroduced to the now familiar context for two consecutive 5-minute sessions separated by a 20-minute interval. During the first session, they were exposed to two identical copies of a novel object (e.g., object A), and during the second session, to two copies of a different novel object (e.g., object B). Animals could freely explore the objects.
- Practice: Sixty minutes after the last encoding session, animals underwent three 3-minute practice sessions separated by 15-minute intervals. The identity of the objects presented depended on the assigned experimental condition (RP, IC, or TC), described below.

RP: One of the two encoded objects was pseudorandomly assigned as the retrieval practice object (RP+). Across three practice sessions, the RP+ object was co-presented with a completely novel object (X, Y, and Z, respectively, in each session). The location (left/right) of each object was randomly assigned per session. This design differs from the original paradigm (Bekinschtein et al., 2018), in which the novel objects X/Y/Z had been pre-exposed in a different context, a manipulation that engages 5-HT2AR-dependent Object-in-Context recognition (Bekinschtein et al., 2013). By using entirely novel objects during retrieval practice, we ensured that any pharmacological effect could be attributed to RIF specifically, rather than to impaired recognition of object-context associations.
IC: This condition allowed us to determine whether the mere presentation and encoding of objects across the three phases was sufficient to induce forgetting through interference. Across the three sessions, animals were exposed in each session to a different pair of identical, completely novel objects (XX, YY, and ZZ).
TC: This time control allowed us to equate forgetting due to the passage of time, in the absence of either retrieval practice or interference from novel object encoding sessions. To achieve this, animals simply remained in their usual cages the entire time, after the initial encoding phase.

Day 3 - Test: Twenty-four hours after the practice phase (or the equivalent time point in the TC condition), we assessed the memories of the two originally studied objects. In the RP condition, we first evaluated the memory for the competitor object by exposing the animals to a copy of the RP-object alongside a completely novel object (object C). Twenty minutes later, we reintroduced the animals to the same context, introducing a copy of the RP+ object and another novel object (object D). We consistently tested memory for the RP-object before the RP+ object to minimize any interference effects that retrieving a stronger memory might have on the attempt to recover a weaker one. In the control conditions (IC and TC), the object whose memory was assessed was randomized across animals. In all cases, the left/right location of the familiar object was counterbalanced across animals.

For the experiment described in Figure 2A, we used a modified version of the task as previously reported (Gallo et al. 2022). Briefly, we increased the number of sessions during the encoding phase, increased the time between the encoding phases and delayed the onset of retrieval practice to 24h.

### Quantification of behavior

Behavior was recorded for each phase using cameras placed over the arenas. Exploration times for each object were then quantified using digital stopwatches (Stopwatch, Center for Behavioral Neuroscience, Emory University, Atlanta, GA). An animal was considered to be exploring an object if it directed its nose toward the object at a distance of 1 cm or less and/or touched the object with its nose. Walking on the object, sitting on it, or using it for support was not considered part of exploration (Ennaceur and Meliani, 1992). Quantification was done blindly by the experimenters. For each phase of the experiment, we measured the total exploration time devoted to each object and derived a preference or discrimination index (DI) based on the relative exploration of the two objects presented. During encoding, this index reflected preference between objects A and B. During practice, it reflected discrimination between object A and the novel objects presented across sessions (X/Y/Z). During the test, it reflected discrimination between the familiar and novel objects. Positive values indicate greater exploration of the preferred or novel object. All raw behavioral data and analysis files used in this study are publicly available in the Open Science Framework (OSF) repository: https://osf.io/5tkqr/overview?view_only=5aacfc99c99549b5b07f121aa7e0dbf1.

### Apparatus

For each condition, animals were exposed to arenas with different visual cues (figures or patterns) on their walls made with self-adhesive paper of different colors. Arena 1 (50 cm width, 50 cm length, 40 cm height) is square-shaped and made of white acrylic. Arena 2 (60 cm width, 40 cm length, 40 cm height) is rectangular and constructed from acrylic; the floor and three walls’ backgrounds are white, while one wall is transparent, allowing unobstructed viewing from one side. Arena 3 (53 cm side, 53 cm height) is triangular, with floor and walls made of white acrylic, except for one wall in contrasting black. Arena 4 (57 cm diameter, 42 cm height) is circular and made of flexible plastic; the floor is white acrylic, while the walls are burgundy.

The assignment of the contexts was randomized; no context was associated with a particular condition or pharmacological treatment across animals. During the animals’ exposure to the context, the lighting of the behavioral room was dim (40 lux) and constant within the contexts (no shadows were generated that could guide the animal’s preference for a particular region of the arena).

### Objects

For all experiments, we used objects that differed in shape, texture, size (between 8 and 24 cm), and color, and that did not have systematically aversive or attractive characteristics to the animals. We had duplicates of all objects to be presented in pairs. Objects varied between weeks of the same experiment for a particular subject and between experiments. The position of the novel object was balanced between treatments, such that novelty was not associated with one specific context position. Objects were attached to the context floor with a reusable odorless adhesive (Patafix, UHU). After each exposure, the context’s objects, floor, and walls were cleaned with 50% ethanol. For clarity, we refer to the objects in the description and diagram of the behavioral task using letters (e.g., object A, object B, etc.). These letters do not indicate that the same objects were used across treatments or between trials. To control for object-related biases, object order was counterbalanced across conditions. In the control conditions (IC and TC), we counterbalanced whether the object evaluated during the test phase had been presented first or second during the encoding phase. In the RP condition, we counterbalanced whether the RP+ object had been presented first or second during the encoding phase.

### Surgery

Guide cannulas (22G) were implanted in the mPFC of both hemispheres, following the coordinates measured relative to the bregma of the skull (anteroposterior +3.20 mm; lateral ±0.75 mm; dorsoventral −3.50 mm; (Paxinos and Watson, 2006)).

After being handled for at least two consecutive days, the rats were habituated to the operating room for at least one hour on the day of surgery. Pharmacological anesthesia was administered intraperitoneally (ketamine Dr. Gray SACI, 80 mg/kg, and xylazine Richmond, 8 mg/kg), and analgesic treatment was administered subcutaneously (meloxicam, 0.2 mg/kg). The effect of anesthesia was confirmed by evaluating the paw reflex to a pinch. The animal was then placed in a stereotaxic frame. Within a time window of 35 to 45 minutes, the necessary craniotomies and cannula implantations were performed. The cannulas were secured to the skull with dental acrylic. At the end of the surgery, animals were injected with a single dose of antibiotic (gentamicin, 0.6 mg/kg) and given a nutritional supplement to prevent weight loss. Handling of the animals resumed at least five days after surgery, after evaluating the absence of signs of distress (e.g., lack of postural reflex or rapid weight loss).

### Cannulae placement

Once the experiments were completed, post-mortem histological controls of the cannula placement were performed. A 1 μl solution of 4% methylene blue in saline was infused through each cannula, and 10-15 minutes later, the animals were sacrificed in CO_2_ chambers following the institutional protocols. The brains were sectioned, and the cannula locations were verified through examination.

### Drug infusion

Handling was done gently to minimize stress. At the time of drug administration, the infusion cannula, with a 0.3 mm gauge (30G), was inserted into the guide cannula. The infusion cannula was 1 mm longer than the guide cannula. After each infusion, the infusion cannula was kept inside the guide cannula for one minute to allow for drug diffusion and prevent reflux.

1μL per hemisphere was infused 10 to 15 minutes before the practice phase of the following solutions or their respective vehicles, following the doses used in previous studies. 8-OH-DPAT: Sigma, 300 ng/µL in saline (Winstanley et al., 2003; Bekinschtein et al., 2013); MDL11,939: Tocris Bioscience, 300 ng/µL DMSO 5% (Carli et al., 2006; Bekinschtein et al., 2013); SB242084: Tocris Bioscience, 300 ng/µ in saline (Takahashi et al., 2001; Bekinschtein et al., 2013); LY294002: Tocris Bioscience, 0.5nmol/µL in 20% DMSO (Fukumoto et al., 2018); PP2: Tocris Bioscience; 33,3 ng/µL in 5% DMSO (Xie et al., 2013); TCB-2: Tocris Bioscience, 4µg/µl in 5% DMSO (Gao et al., 2020); WAY 100135: Tocris Bioscience, 2 µg/µl in 5% DMSO (Carli et al., 1995).

### Immunoblot assays

Animals underwent surgery as previously described and, after recovery, were assigned to either the IC or RP condition. Within each condition, animals were further divided according to pharmacological treatment (VEH or LY) and then subjected to the RIF protocol, including the sample and practice phases. Rats were sacrificed by decapitation immediately after the second practice session. The mPFC was rapidly dissected by the experimenter on an ice-cold plate within 3 min, and the tissue was then stored at −80 °C until homogenization. Tissue was homogenized in ice-chilled buffer (pH 7.4, 20 mM Tris-HCl, 0.32 M sucrose, 1 mM EDTA, 1 mM EGTA, 1 mM PMSF, 10 μg/ml aprotinin, 15 μg/ml leupeptin, 10 μg/ml bacitracin, 10 μg/ml pepstatin, 15 μg/ml trypsin inhibitor, 50 mM NaF, 1 mM sodium orthovanadate and 0.1 mM ammonium molybdate) (de Landeta et al., 2020) and samples were stored at −80 °C until used. Samples (7.5 μg of proteins, determinated by Pierce BCA Protein Assay Kit, Thermo Scientific USA) were subjected to SDS-PAGE (polyacrylamide 10%). Proteins were transferred to Polyvinylidene fluoride (PVDF) membranes overnight at 4 °C. Membranes were incubated first with anti-phospho Thr 308 AKT antibody ((D25E6) 1:2000, Rabbit monoclonal, Cell Signalling, USA), then stripped and incubated with anti-total AKT antibody (9272= Antigen-antibody complexes were visualized by a fluorescent method using ECF substrate (GE Healthcare Life Sciences, USA) and fluorescence-measuring equipment (STORM scanner, GE Healthcare Life Sciences, USA). Densitometry analysis was performed using Gel-Pro Analyzer (version 4.0 Media Cybernetics, USA), optical density (OD) values were obtained for each sample for both p-AKT and total AKT. Each p-AKT OD value was normalized to its corresponding total AKT OD value, and then normalized to IC-Vehicle group. Planned pairwise contrasts were performed to examine simple effects within the interaction. Given the small sample size and the a priori nature of these comparisons, p-values were not adjusted for multiple comparisons.

### Experimental design and statistical analysis

Mixed experimental designs were used for all the experiments. The animals were randomly assigned to one of the three experimental conditions (RP, IC, or TC) and, in different weeks, received the possible pharmacological treatments (i.e., drug and vehicle). The sample size used was based on results from previous studies in the laboratory, assuming that if a treatment effect were found, it would be of a similar magnitude (Bekinschtein et al., 2018; Gallo et al., 2022).

Linear mixed models were used, detailed in each case, to determine the statistical significance of fixed effects, taking into account possible interactions between factors. Subject was included as a random effect to account for repeated measures. Post-hoc contrasts (Bonferroni) were performed, adjusting the p-value for multiple comparisons. Memory expression was additionally assessed by one-sample t-tests against zero.

To assess whether initial object preference during encoding systematically influenced test performance, we examined the Pearson correlation between encoding exploration preference and the test Discrimination Index across experiments and control conditions. Although isolated significant correlations emerged in individual groups, no consistent pattern was observed across experiments, suggesting that encoding bias did not systematically confound test performance. We verified the assumptions of normality and homoscedasticity of the residuals for the primary outcome measures using the Shapiro-Wilk and Levene tests, respectively.

A significance level of 0.05 was used throughout. All data processing and statistical analysis were conducted in R using RStudio (version 2023.06.1). Linear mixed models were fit with the nlme package, post-hoc contrasts were performed with emmeans, and Levene tests with car.

#### Subject exclusion criteria

We initially included 268 subjects. Based on the criteria detailed below, 25 were excluded, resulting in a final sample of 243 subjects.

Behavioral: exclusion criteria were established based on the animals’ exploratory behavior over the different phases of the protocol. Animals were excluded if they did not meet all of the following requirements:

Encoding: total exploration time greater than 10 s per encoding session.
Practice (RP): total exploration time for the familiar and novel objects greater than 10 s across practice sessions.
Practice (IC): exploration time greater than 5 s in each practice session.
Test: total exploration time greater than 10 s.
Statistical: extreme values (±3 standard deviations from the mean) were identified for all calculated variables and excluded from the analysis.
Other: subjects who showed signs of distress (e.g., due to the cannula implantation surgery) were excluded and sacrificed immediately. Animals were also excluded if they learned to escape from the context or had incorrect cannula placement.

## Acknowledgment

This work was funded by by the Consejo Nacional de Investigaciones Científicas y Técnicas, PUE 0052; Fondo para la Investigación Científica y Tecnológica [FONCyT; Grants PICT 2018-1063 (NW), PICT 2019-2544 (NW), PICT 2018-1062 (PB), PICT 2021-4663. Medical Research Council grant MC-A060-5PR00 (MCA). During the preparation of this work the authors used OpenAI’s ChatGPT (April 2024 version) and Anthropic’s Claude (Opus 4.7, August 2026) to improve clarity and style. After using these tools, the authors reviewed and edited the content as needed and take full responsibility for the content of the publication. We thank Dr. Victoria Oberholzer for helpful discussions on the manuscript, Jaime David, and Vet. Maria Ines Bensansón for technical assistance.

## Author contributions

MBZ, PB, and NW designed the experiments. MBZ, MI, and PK collected the behavioral data. BAO, MB, PK and CK performed the Western blot experiments. MBZ, PB, MC, and NW wrote the manuscript. All authors reviewed the manuscript and approved the final version.

